# An emerging target paradigm evokes fast visuomotor responses on human upper limb muscles

**DOI:** 10.1101/2020.04.07.030130

**Authors:** Rebecca A. Kozak, Aaron L. Cecala, Brian D. Corneil

**Affiliations:** Graduate Program in Neuroscience, Western University, London, On, Canada, N6A 5B7; Department of Physiology and Pharmacology, Western University, London, Ontario, Canada, N6A 5B7; Department of Psychology, Western University, London, Ontario, Canada, N6A 5B7; Robarts Research Institute, 1151 Richmond St. N, London, Ontario, Canada, N6A 5B7

**Keywords:** Stimulus-locked responses (SLRs), Reaction time (RT), Visually-guided reaches, Humans, Electromyography

## Abstract

To reach towards a seen object, visual information has to be transformed into motor commands. Visual information such as the object’s colour, shape, and size is processed and integrated within numerous brain areas, then ultimately relayed to the motor periphery. In some instances we must react as fast as possible. These fast visuomotor transformations, and their underlying neurological substrates, are poorly understood in humans as they have lacked a reliable biomarker. Stimulus-locked responses (SLRs) are short latency (<100 ms) bursts of electromyographic (EMG) activity representing the first wave of muscle recruitment influenced by visual stimulus presentation. SLRs provide a quantifiable output of rapid visuomotor transformations, but SLRs have not been consistently observed in all subjects in past studies. Here we describe a new, behavioural paradigm featuring the sudden emergence of a moving target below an obstacle that consistently evokes robust SLRs. Human participants generated visually-guided reaches toward or away from the emerging target using a robotic manipulandum while surface electrodes recorded EMG activity from the pectoralis major muscle. In comparison to previous studies that investigated SLRs using static stimuli, the SLRs evoked with this emerging target paradigm were larger, evolved earlier, and were present in all participants. Reach reaction times (RTs) were also expedited in the emerging target paradigm. This paradigm affords numerous opportunities for modification that could permit systematic study of the impact of various sensory, cognitive, and motor manipulations on fast visuomotor responses. Overall, our results demonstrate that an emerging target paradigm is capable of consistently and robustly evoking activity within a fast visuomotor system.

**SUMMARY:** We present a new behavioual paradigm that elicits robust fast visuomotor responses on human upper limb muscles during visually guided reaches.

## INTRODUCTION

When we notice a message on our cellphone, we are prompted to perform a visually-guided reach to pick up our phone and read the message. Visual features such as the shape and size of the phone are transformed into motor commands allowing us to successfully reach our goal. Such visuomotor transformations may be studied in laboratory conditions, which permit a high degree of control. However, there are scenarios where response time is of the essence; for example, catching our phone if it were to start to fall. Laboratory studies of fast visuomotor behaviors often rely on displaced target paradigms where on-going movements are modified in mid-flight following some change in target position (e.g.-(Soechting and Lacquaniti, 1983; Veerman et al., 2008)). While such online corrections can occur in <150 ms (Day and Lyon, 2000), it is difficult to ascertain the exact timing of fast visuomotor output using kinematics alone due to the low-pass filtering characteristics of the arm, and because such output is superseding a movement already in mid-flight. Such complications lead to uncertainty about the substrates underlying fast visuomotor responses (e.g.-see review by(Gaveau et al., 2014)). Some studies suggest that subcortical structures such as the superior colliculus, rather than fronto-parietal cortical areas, may initiate online correction (Day and Brown, 2001).

This uncertainty regarding underlying neural substrates may be due, at least in part, to the lack of a reliable biomarker for the output of the fast visuomotor system. Recently, we have described a novel measure of fast visuomotor responses that may be generated from static postures and recorded via electromyography (EMG). Stimulus-locked responses (SLRs) are time locked bursts of EMG activity that precede voluntary movement (Corneil et al., 2004; Pruszynski et al., 2010), evolving consistently ∼100 ms after stimulus onset. As the name implies, SLRs are evoked by stimulus onset, persisting even if an eventual movement is withheld (Wood et al., 2015) or moves in the opposite direction (Gu et al., 2016). Furthermore, SLRs evoked by target displacement in a dynamic paradigm are associated with shorter latency online corrections, and manipulations of sensory input that influence SLR timing also influence the timing of on-line reach corrections (Kozak et al., 2019). Thus, SLRs provide an objective measure to systematically study the output of a fast visuomotor system, as they may be generated from a static posture and parsed from other EMG signals unrelated to the initial phase of the fast visuomotor response.

All studies investigating the SLR have reported less than 100% detection rates across participants, even when using more invasive intramuscular recordings (Pruszynski et al., 2010; Wood et al., 2015; Gu et al., 2016). Low detection rates and a reliance on invasive recordings limit the usefulness SLR measures in future investigations into the fast visuomotor system in disease or across the lifespan. While it is possible that all subjects may simply not express the SLR, it may also be the case that the stimuli and behavioural paradigm were not ideal. Past reports of SLRs have used paradigms wherein participants generate visually-guided reaches towards static, suddenly appearing targets (Pruszynski et al., 2010; Gu et al., 2016). However, a fast visuomotor system is most likely needed in scenarios where we must rapidly interact with a falling or flying object, leading us to wonder if moving rather than static stimuli may better evoke SLRs. Here, we have adapted a moving target paradigm used to study eye movements (Kowler, 1989), and combined it with a pro/anti visually guided reaching task used to examine the SLR (Gu et al., 2016). We found the emerging target paradigm improved the magnitude and detectability of the SLR, in comparison to past studies relying on static stimuli. The emerging target paradigm also elicited shorter latency RTs. The purpose of this paper is to describe the emerging target paradigm, and present results on SLR magnitude, across-subject prevalence and RT in comparison to the static paradigm used previously. The emerging target paradigm can also be modified in many different ways, which should promote more thorough investigations into sensory, cognitive, or motor factors that promote or modify fast visuomotor responses.

## PROTOCOL

### 1. Participant preparation

1.1 After obtaining informed consent, participant preparation involves the application of EMGs to the upper limb muscles and setup in the apparatus. The pectoralis major muscle is involved in cross body reaching and provides a convenient location for electrode placement; therefore, the remainder of the methods will focus on this muscle. You may target the clavicular head, and if this location is not ideal then the sternal head, or the lateral portion of the pectoralis muscle close to the shoulder. Reasons for alternative electrode placement may include excessive hair or distribution of adipose, both of which may lower the signal to noise ratio of the EMG signal. To ensure consistency within this experiment, we required all participants to use their right arm for visually guided reaches regardless of handedness. Here we used a commercially available system to record surface EMG activity (Delsys Inc. Bagnoli-8 system), however other recording equipment or electrodes may be used.

> 1.1.1 Visualize the muscles from which you will be recording. For pectoralis major, ask participants to relax elbows at sides and push palms together. This action will allow the visualization of target muscle(s), as this action is similar to the one used to grasp and manipulate the manipulandum.
>
> 1.1.2 Using alcohol swabs, clean the skin surface overlying the sternal and/or clavicular heads of the pectoralis major of the reaching arm and skin overlying the contralateral clavical; the latter will be the location of the ground electrode.
>
> 1.1.3 Apply double sided adhesive to Delsys surface sensors then apply small amount of electrode gel to each of the parallel EMG sensors.
>
> 1.1.4 Ask participant to place palms together again and adhere sensors perpendicular to the long axis of the visualized target muscle. Place ground electrode on left clavicle. Secure sensors and ground electrodes to surrounding skin with medical tape and insert sensor wires into Delsys amplifier.
>
> 1.1.5 We used a desktop computer connected to the Delsys system to provide real-time monitoring of EMG activity throughout the experiment, and to help determine the suitability of electrode placement. To determine a suitable placement, ask participant to fully extend their right arm. During cross-body, leftward reaches, the peak EMG activity should be at least 2 times the level of baseline activity (if not considerably higher), whereas EMG activity should decrease from baseline during movements in the opposite (right) direction. Reposition the electrodes if necessary to ensure that these activity levels are observed.

1.2 We have set up participants in a bilateral endpoint robot (Kinarm, Kingston, Ontario, Canada). Only the right manipulandum was used. SLRs have been reported in the context of upper limb movements made in other platforms. The Kinarm endpoint robot allows reaching movements in a horizontal plane, and the application of force to the manipulandum. Adding force increases the background activity of the muscle of interest, allowing for expression of the SLR as an increase or decrease in muscle activity following stimulus presentation in the muscle’s preferred or non-preferred direction, respectively. A level of baseline activity is especially useful in the non-preferred direction, as baseline and non-preferred reaching activity in the pectoralis muscle would be indistinguishable without a background loading force. We have applied 5N of force to the right and 2N of force down (opposite to a leftward presented target relative to the start position), throughout the entirety of the experiment.

> 1.2.1 Seat the participant in the experimental chair. Move and adjust the chair close enough to the screen and robot so the participant is comfortable. With a constant force applied to the arm for the duration of the experiment, the priority is to ensure the participant is comfortable, as repeated changes in body posture may change background muscle recruitment, while also ensuring that the recorded muscle is recruited during the reaching task.

### 2. Stimuli construction/ apparatus

2.1 All experimental procedures used the KINARM endpoint robot system. All stimuli were generated via the Kinarm apparatus using Matlab® (version R2016a, The MathWorks, Inc., Natick, Massachusetts, United States) Stateflow® and Simulink® applications. In our setup, stimuli were presented via a VPIXX projector (Saint-Bruno, QC, Canada) custom integrated into the Kinarm platform to ensure high quality visual images and event timing.

> 2.1.1 The emerging target paradigm contains 4 primary components, all of which are referenced here in relation to the midpoint of the two robot manipulandum origins in the Kinarm endpoint robot (reported in cm). An **inverted y path** (y: - 19 (top of inverted y) or -34 (bottom of inverted y), x:-/+2 (inner, bottom inverted y), -/+8 (outer bottom inverted y); width .5 height: 20 (top) or 15 (bottom)), an **occluder** (centered at: 0, -29; width: 35 height: 15), a **moving target** which moves down the inverted y and behind the occluder (start: 0, -17; radius: 1; speed: 10 cm/s, speed behind occluder: 30 cm/s), and **start position** (0, -42; radius 1) (See Supplementary material for screen shot).
>
> 2.1.2 The occluder contains a notch cut out on the center bottom between the two outputs (0, -29; width: 5 height: 5). The participant is instructed to: “fixate the notch while a target is behind the occluder”. Doing so ensures the eye is stable at target emergence.

### 3. Procedure

3.1 Throughout a trial, the distal portions of the “inverted Y” path, the occluder box, and a white cursor representing the participant’s hand position were present (See Supplementary material for screen shot). The participant’s hand/arm was occluded during the experiment via an upward facing mirror reflecting downward-presented targets. Hand position was represented in the position as a real-time cursor (RTC) projected onto the screen. At the beginning of each trial, two stationary white dots are also presented to the particpant. The dot above the occluder (T1) will become the eventual reach target. The other dot (T0) was located below the occluder, and represents the central start position.

> 3.1.1 For each trial, a participant brings the RTC into the central start position T0. Motion of T1 starts once the RTC remains aligned with T0 for a variable duration of 1-1.5 s. If the participant exits the T0 start position before the prescribed time, the trial will start again. The occluder will either be green, instructing the subject to generate a pro-reach toward the target when it emerges from below the occluder, or red, instructing the subject to generate an anti-reach away from the emerging target. T0 disappeared once T1 begins moving, at which point no restrictions are placed on the subject’s arm motion.
>
> 3.1.2 After T1 reaches the occluder, it moves behind the occluder and travels at a constant speed of 30 cm/s along the y axis. Once T1 reaches half the length of the occluder, it bifurcates along one of the inverted y outputs with an additional x axis speed of 30cm/s. Thus, speed along the y axis is kept constant. The side T1 appears at the bottom of the inverted ‘y’ is randomized. The target vanishes for ∼0.5 s, depending on the size of the occluder and the speed of T1 motion.
>
> 3.1.3 Once T1 reaches the edge of the occluder closest to the participant, instead of sliding past the edge of the occluder to create an initial ‘half-moon’ stimulus, T1 is invisible to the participant until the full target has moved below the occluder. This is done to control for visual processing effects of partial stimuli, especially if different speeds of targets are used which would cross the boundary at different times. A partial emergence of a target (e.g.-half moon stimulus) produce a target composed initially of a higher spatial frequency, which based on our previous results would lead to increased SLR latency and decreased magnitude (Kozak et al., 2019).
>
> 3.1.4 Simultaneous with target appearance, a secondary target is presented in the corner of the screen, at a location covered by a photodiode. This target presented to the “corner” photodiode is not seen by the subject, but provides an analog signal that allows for the precise alignment of target appearance with muscle activity. This veridical signal is needed to account for any delay between the time that presentation software requests that the target emerges below the obstacle, and the time at which such emergence actually happens. Prior to data collection in any setup, we recommend using multiple photodiodes (one over the corner stimulus, others over where the targets emerge) to identify any lags in stimulus timing and ensure minimal trial-by-trial variability.
>
> 3.1.5 If the occluder is green, the participant is instructed to perform a pro-reach towards the emerging target, and intersect it, at which point the trial will end. If the occluder is red the participant must perform an anti-reach in the opposite direction of the target. The ‘inverted y’ path of T1 promotes leftward and rightward reaches in the pectoralis major muscles preferred and non-preferred direction, respectively.
>
> 3.1.6 Correct feedback is provided after the trial as either a ‘HIT’ (correct interception), ‘WRONGWAY’ (incorrect direction for pro/anti reach), or ‘MISS’ (neither correct nor incorrect responses detected). This feedback is written in the middle of the occluder in the inter-trial interval. In the anti-reach conditions, a correct interception (moving away from a target) is not based on the mirror image of T1, but rather the horizontal distance relative to T0. Distance is also used to determine a ‘MISS’ trial, where a target had moved a certain distance off of the screen and no correct or incorrect response was detected.

3.2 The procedure was split into 4 blocks of 100 trials each. Trial types were randomly intermixed. Each participant performed 100 reaches per unique condition (targets emerging left or right, pro- or anti-reach conditions; 4 trial types total). It is recommended that each condition consists of a minimum of ∼80 reaches per condition when using surface recordings, as the next analysis step relies on data from many trials for SLR detection. Each block took approximately 7.5 minutes to complete.

3.3 For the purpose of comparison, we have also included a ‘static’ pro-reach and anti-reach paradigm, similar to paradigm reported in (Gu et al., 2016). Doing so allows us to compare the properties of SLRs recorded in the emerging target task to those recorded in a task used previously. Briefly, all recording methods are identical as listed above. The key difference lies in the nature of the behavioural paradigm, which involves only T0 (at the start position) and the presentation of a static target to the left or right. The start position (T0) is in the same position as in the emerging target paradigm (y: 0, x: -42, relative to the midpoint between the robotic manipulandum origins). The radius of T0 is larger (r=2), and the colour of T0 changes to red or green to instruct the subject to generation a pro or anti reach in response to T1. T target (T1) appears 10 cm to the left or right of T0 after a randomized hold period (1-2s where the participants RTC must remain inside of T0; this period also serves as the instruction period based on the color of T0). As in the emerging target paradigm, a participant must reach towards a target if T0 is green, and reach in the diametrically opposite direction away from a target if T0 is red. As in Gu et al. 2016, the appearance of T1 was synchronous with the disappearance of T0.

### 4. Analysis

Each block of data gathered from a participant has a downloadable .zip file containing kinematic data from each trial, as well as Delsys EMG surface recordings and photodiode output saved as analog inputs. These files are then unzipped and analyzed offline via custom Matlab scripts. Data was sampled at 1000 Hz. Error trials were not analyzed, as indicated by incorrect reach directions (3.5 cm in the wrong direction), long RTs (>500 ms) indicating presumed inattentiveness or short RTs (<120) indicating anticipation. The following details our method of analysis of SLRs and RTs, although alternative methods may also be suitable.

> 4.1.1 Reaction time was defined as 8% of the peak tangential velocity.
>
> 4.1.2 To analyze muscle activity offline, customized Matlab scripts convert the signals to source microvolts, and remove any DC offset, rectify the EMG signal, and filter the signal with a 7-point moving average filter.
>
> 4.1.3 A time-series receiver-operating characteristic (ROC) analysis is used to detect the presence and latency of the SLR (Corneil et al., 2004; Pruszynski et al., 2010), by defining the probability by which an ideal observer can discriminate through time the side of stimulus presentation based only on EMG activity. In our experiment, the ROC values were based on two distributions of EMG activity following leftward or rightward stimulus presentation (e.g.-Fig. 2c red versus light red traces). The ROC value (i.e., the area under the ROC curve) was calculated and plotted at every time sample (1ms) from 100 ms before to 300 ms after target presentation (Fig. 1b or 2c). A ROC value of .5 indicates chance discrimination, whereas values of 1 or 0 indicate perfectly correct or incorrect discrimination relative to target presentation, respectively. Discrimination thresholds was set to 0.6, and the discrimination latency was determined as the first of 8 of 10 consecutive points that exceeded this value (Fig. 2c indicated by vertical red or blue lines). These parameters (threshold, and number of points exceeding threshold) may change depending on the quality and quantity of surface or intramuscular EMG recordings, and a bootstrapping analysis may be used to objectively determine confidence intervals. Past work of ours has shown that a value of 0.6 equates approximately to a 95% confidence interval (Goonetilleke et al., 2015).

**FIGURE 1.**
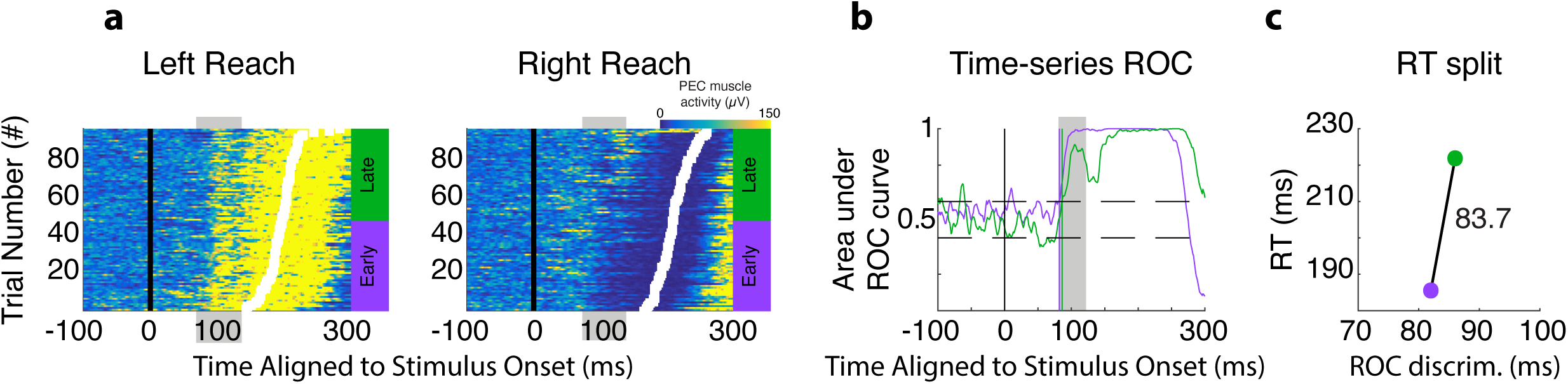
Example of an SLR from a representative participant, illustrating our detection criteria. a) Trial-by-trial recruitment for right pectoralis major muscle for right or left reaches in the pro-reach condition. Each row is a different trial. Intensity of colour conveys the magnitude of EMG activity. Trials are sorted by reach RT (white boxes) and aligned to stimulus onset (black line). The SLR appears as a vertical banding of activity highlighted by grey boxes; note how EMG activity increases or decreases time-locked ∼90 ms after leftward or rightward stimulus presentation, respectively. Purple or green bars indicate the trials contributing to the early or late RT groups, respectively. b) Time-series ROC analysis indicating time of EMG discrimination for early (purple) and late (green) trials shown in (a). c) For the early (purple) and late (green) gropus, mean RT is plotted as a function of ROC discrimination. The slope of the line connecting these two points is 83.7 degrees, indicating that EMG activity is more aligned to stimulus presentation than movement onset. SLR detection requires that the slope of this line exceed 67.5 degrees (halfway between 45 and 90).

**FIGURE 2.**
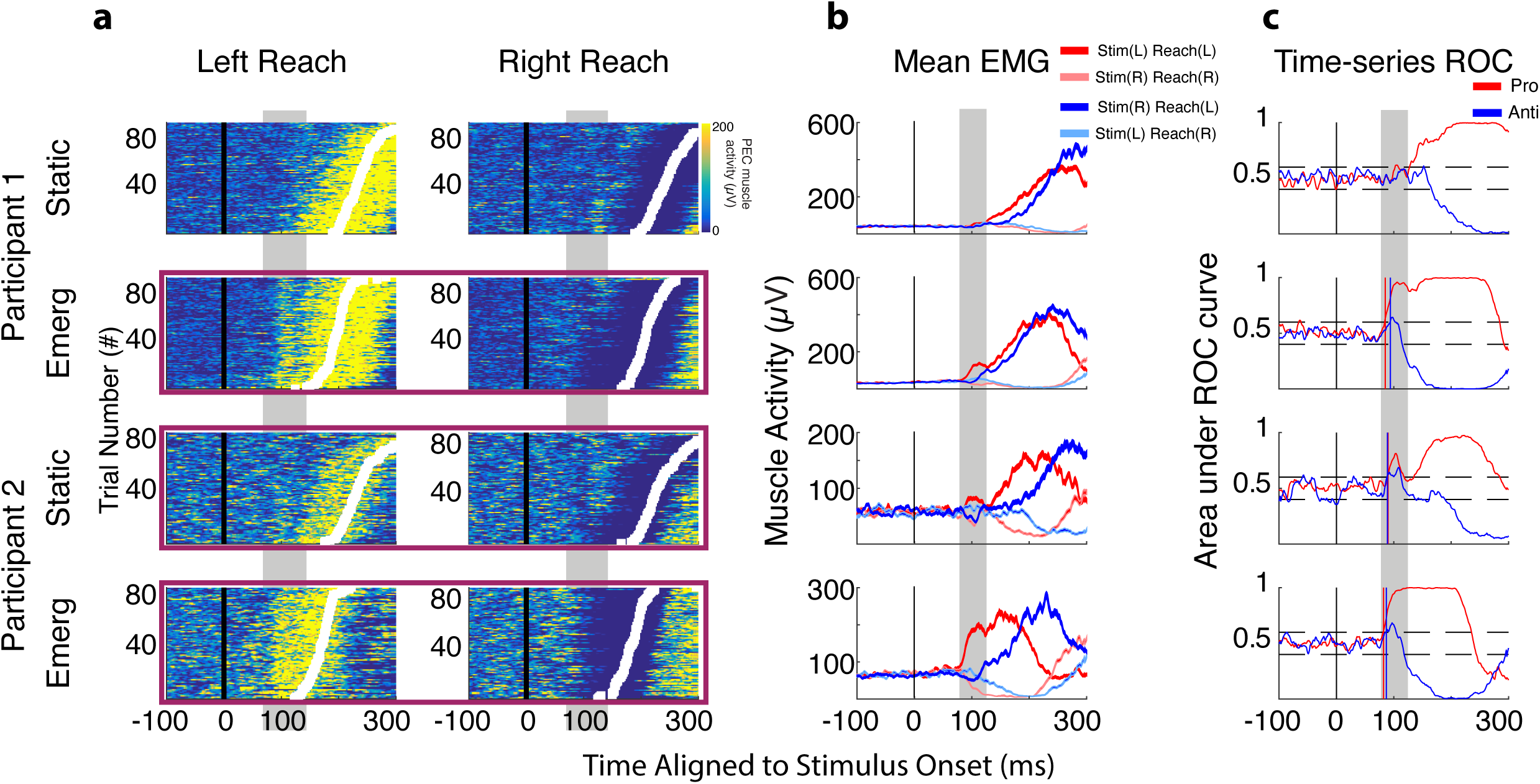
Representative results from participants 1 and 2, showing the variability in the presence or absence or SLRs in the static (1^st^ and 3^rd^ rows), and the consistency of SLR presence in the emerging target paradigms (2^nd^ and 4^th^ rows). A) Trial-by-trial recruitment for right pectoralis major muscle for these participants (same format as Fig. 1a). Conditions exhibiting an SLR are outlined in purple (2^nd^, 3^rd^ and 4^th^ rows). B) Mean +/- SE of EMG activity for both pro (red) and anti (blue) reaches, segregated by side of stimulus presentation (fainter traces used for movements in the non-preferred direction). Note how EMG activity in the SLR interval often initially increases after leftward stimulus presentation, even on anti-reach trials where the reaches proceed to the right (light blue traces). C) Time-series ROC analysis for pro (red) and anti (blue) reaches shown in (b). SLR epoch highlighted in grey box; horizontal dashed lines at 0.4 and 0.6. Vertical coloured lines (if present in pro condition) show the discrimination time for pro- (red) or anti- (blue) reach trials.

> 4.1.3.1 We then determined the **presence** of an SLR on pro-reach trials using a RT-split analysis (see Fig.1, (Wood et al., 2015)). Briefly, this analysis determines whether EMG activity is more locked to stimulus onset rather than movement onset by using two separate time-series ROC analyses. First, trials from left and right directions are sorted and divided in ‘early’ and ‘late’ RT groups (Fig. 1a). A time-series ROC analysis is performed separately on early and late groups (Fig. 1b), which yields separate EMG discrimination times for both groups. The ROC discrimination times are then plotted against the mean RTs for their respective early and late groups. This plot yields two points (early discrimination/RT versus late discrimination/RT) that are connected via a line (Fig. 1c). For this line, a slope of 90 deg would indicate that EMG discrimination times are locked to stimulus presentation, whereas a slope of 45 deg would indicate that EMG discrimination are locked to movement onset. In practice, we used a cut-off slope of 67.5 degrees (halfway between 45 and 90 deg) to detect whether an SLR was present (slope > 67.5 deg) or not (slope < 67.5 deg).
>
> 4.1.3.2 If SLR presence is determined, the **SLR latency** is defined by the discrimination latency from all the trials (4.1.3). After finding the discrimination latency, the same opposing left and right reaches were used to determine the **SLR magnitude** of the response. Left and right mean EMG traces were overlaid on the same plot (Fig. 2c dark red versus light red traces). Magnitude is calculated as the difference between left and right mean EMG traces from SLR latency to 30 ms post discrimination latency.

## TABLE OF MATERIALS

TABLE OF MATERIALS:
| Name of Material/ Equipment | Company | Catalog Number | Comments/Description |
| --- | --- | --- | --- |
| **Kinarm End-Point Robot** | Kinarm |  | Other reaching apparatus may be used |
| **Bagnoli Desktop System** | Delsys |  | Other EMG recording equipment may be used. |

## SUPPLEMENTARY FILES

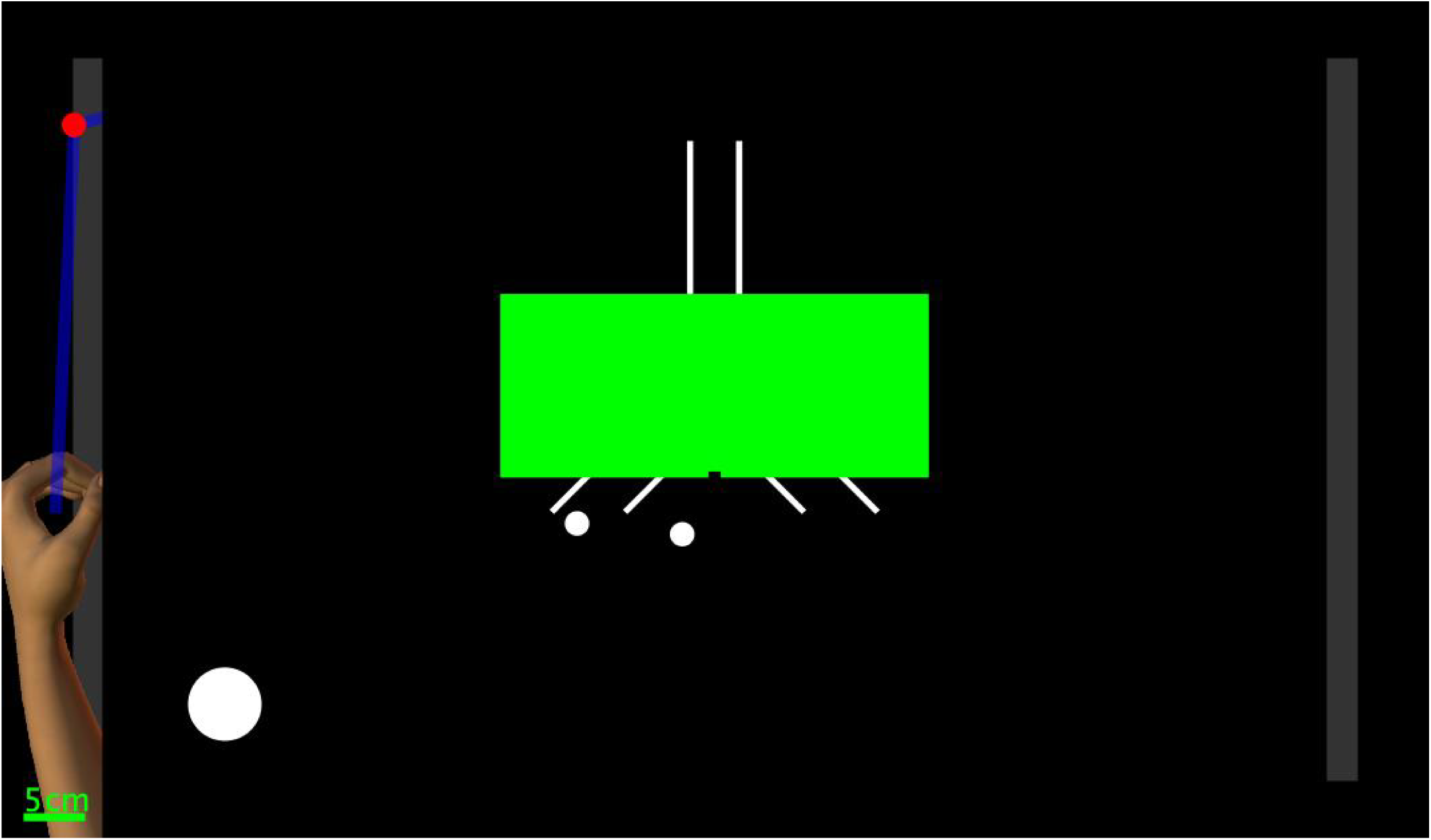
Supplementary Fig. 1.

## DISCUSSION

Humans have a remarkably capacity, when needed, to generate rapid, visually-guided actions at latencies that approach minimal afferent and efferent conduction delays. Systematic study of the neural mechanisms underlying rapid, visually-guided reaches is complicated by the arm’s inertia and the fact that responses such as rapid on-line corrections supersede a movement already in mid-flight. We have previously described stimulus-locked responses (SLRs) on the upper limb as a new measure for rapid visuomotor responses (Pruszynski et al., 2010; Gu et al., 2016; Kozak et al., 2019). While beneficial in providing a trial-by-trial benchmark for the first aspect of upper limb muscle recruitment influenced by the visual stimulus, limb SLRs have not been expressed in all subjects and often relied upon invasive intramuscular recordings. Here, we describe an ‘emerging target paradigm’, in which subjects reach from a stable posture in response to the emergence of a moving visual target from behind an occluder. The benefits of the emergent target paradigm are apparent within individual participants, as participants who does not express the SLR in a paradigm used previously express one in the emerging target paradigm (e.g., Fig. 2, participant 1-1^st^ row versus 2^nd^ row). Furthermore, SLRs expressed in the emerging target paradigm are much larger than in other paradigms, sometimes attaining magnitudes that approach that obtained just before movement onset (Fig. 2, participant 2; Fig. 4, participant 5). Thus, this paradigm has proven to be effective in increasing the magnitude (Fig. 3a), detectability of the SLR (Fig. 3b), and promoting shorter RTs by approximately 50 ms (Fig. 3b), compared to the use of static targets.

**FIGURE 3.**
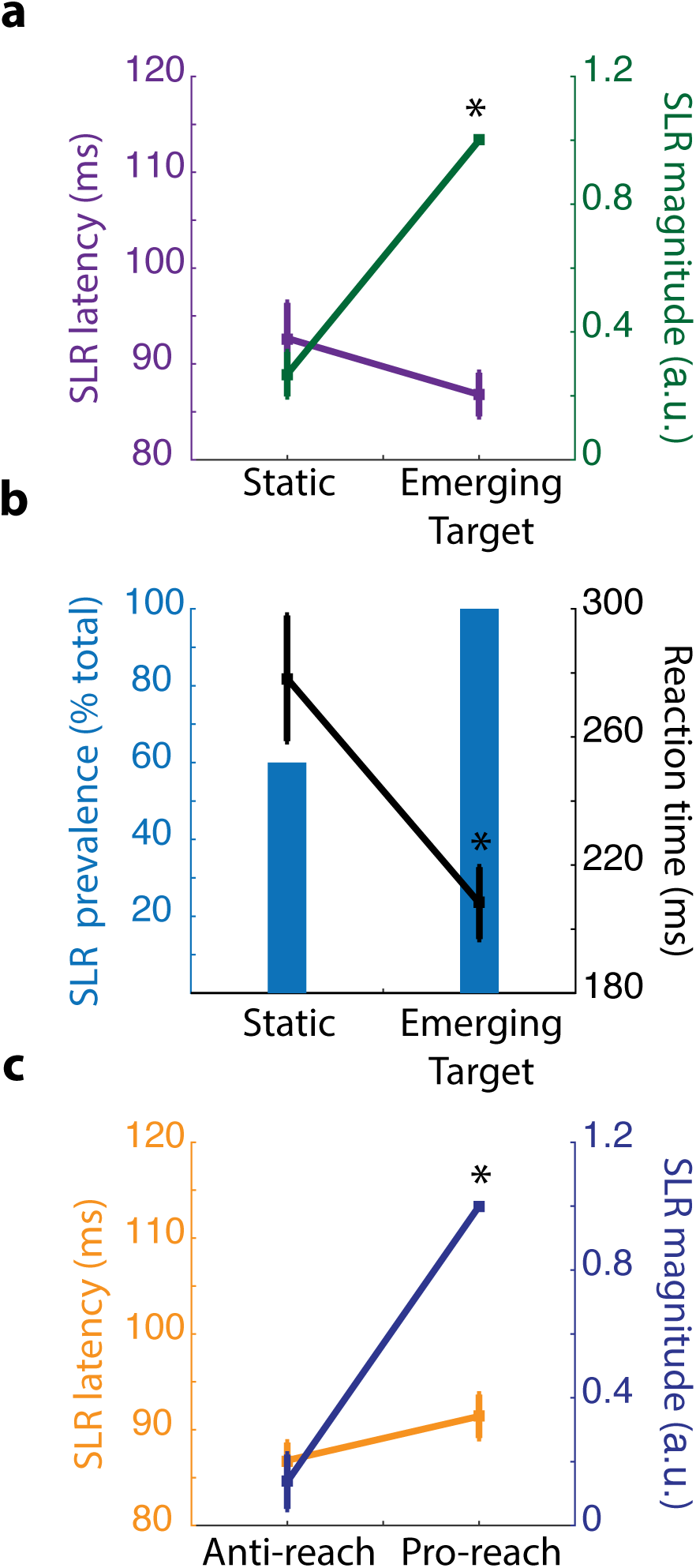
Effects of an emerging target paradigm on SLR characteristics and reach RT. A) SLR latency (purple) and magnitude (green) for pro reaches in static versus emerging target paradigms. Latency defined as first 8 out of 10 continuous data points surpassing ROC threshold of .6 (see methods). Magnitude of SLR is defined as the integrated area over 30 ms after SLR discrimination between the mean EMG activity on left versus right trials. All magnitudes were normalized to the maximum for the participant across conditions (e.g.- a value of 1 indicates the maximal response). B) SLR prevalence and reach RT. SLR prevalence determined with RT split analysis (see methods and Fig. 1). RT determined at 8% of the peak tangential velocity. C) SLR magnitude and latency results from pro and anti-reaches in the emerging target paradigm. A) and B) demonstrates how fast visuomotor responses, be they SLRs or reach RTs, and expedited in the emerging target versus static paradigm. C) shows how a cognitive manipulation to prepare for a pro- or anti-reach influences SLR magnitude but not timing. * denotes significance at p<.05 compared to static or anti condition based on unpaired t-test.

Which features of the emerging target paradigm lead to robust expression of fast visuomotor responses? We speculate that a critical aspect is the implied motion of the target behind the occluder (supplemental material Fig. 1). Implied motion produces strong signals in motion-related areas in the dorsal visual stream that are indistinguishable from those produced by visible moving targets (Krekelberg et al., 2005). Our implementation of the emerging target paradigm also incorporated a high degree of certainty of the time at which the target would re-appear. The disappearance and subsequent emergence of the target behind the barrier may be akin to a ‘gap interval’ between offset of a central fixation or hold stimulus and presentation of a peripheral target, which also expedites reach reaction times (Gribble et al., 2002) and promotes the expression of express saccades (Paré and Munoz, 1996), which are another type of fast visuomotor response. Finally, it is important that the target emerging from behind the barrier is presented in its entirety, rather than being presented as sliding from behind the barrier. Were the target to slide past the barrier, the earliest stimulus available to the visual system would be a ‘half-moon’ stimulus that would lack the lower spatial frequencies known to promote earlier and stronger expression of limb SLRs (Kozak et al., 2019). In addition to these theoretical considerations, it is important to position the outlets for the emerging targets at locations associated with the preferred or non-preferred direction of the muscle(s) under study. Introducing a background loading force to increase activity of the muscle of interest is also beneficial in the detection of limb SLRs, as recruitment would increase or decrease for presentation of the target in the preferred or non-preferred direction, respectively. Finally, given the short latency of the limb SLR, it is imperative to ensure that the time of target emergence is known on every trial; depending on make and model, digital screens or projectors can introduce quite variable delays in stimulus presentation which could compromise accurate alignment of muscle activity to critical events.

There are a number of ways in which the emerging target paradigm could be modified, and doing so can further the understanding of the sensory, cognitive, and movement-related factors that influence the fast visuomotor system. Here, we instructed the subjects to prepare to move toward (a pro-reach) or away (an anti-reach) from the emerging target. As expected from previous results (Gu et al., 2016), consolidation of this instruction enabled subjects to dampen SLR magnitude without changing SLR timing. This shows that the neural centres mediating the SLR can be pre-set by higher-order areas establishing task set, prior to target emergence. There are numerous other dimensions in which the task could be modified to manipulate cognitive factors, for example by altering the predictability of target appearance in either time (i.e., making the timing of emergence less predictable) or space (i.e., biasing target emergence to one side or another, or providing endogenous cues to indicate the side of emergence). Manipulations of the sensory parameters of the emerging target (e.g., the speed, contrast, size, or colour of the emerging stimulus, or the presence of competing distractors) will also provide insights into underlying substrates. Presenting a static rather than moving target below the barrier would also help parse the effects of target motion versus temporal predictability on the robustness of the limb SLR. Finally, from a motor perspective, the framework of the emerging target paradigm can be extended to bilateral reaching movements, and establishing the presence of robust SLRs on upper limb muscles potentiates the investigation the distribution of such signals to other trunk or limb muscles.

One of the challenges associated with this paradigm, perhaps paradoxically, is the degree to which reach RTs were shortened. Our SLR detection criteria resembled that used previously (Goonetilleke et al., 2015), as we ran separate time-series ROC analyses for the shorter- or longer-than median RT groups. Doing so requires some degree of variance in reach RTs, and in practice we have found that reach RTs are shorter and less variable in the emerging target paradigm compared to the static paradigm (279 +/- 58 ms (static); 207 +/- 34 ms (dynamic)). Indeed, RTs were sometimes shortened to such a degree that the movement-related volley of EMG activity often blended into the SLR interval. Consequently, the time-series ROC often rose directly from values near 0.5 to values near 1.0, without displaying the brief decrease after the SLR that was required for detection in ((Wood et al., 2015); see Fig 4, participant 1,2,4,5). We expect that the detection criteria for SLRs may continue to evolve and will likely have to be optimized to the specifics of the task at hand. Other task manipulations, perhaps by increasing the temporal uncertainty of target re-emergence or requiring that subjects wait to move for a short interval after target emergence (e.g., by waiting for the emerged target to change colour), may help increase the mean and variance of reach RTs and separate recruitment during the SLR interval from that associated with movement onset.

In closing, and mindful of the challenges associated with shorter and more invariant reach RTs, the framework of the emerging target paradigm will advance the study of rapid visuomotor responses, by providing a means to obtain robust expression of upper limb SLRs. It is particularly noteworthy that all of the results reported here were obtained with surface recordings, as this will enable study of SLRs in populations that may be less amenable to intramuscular recording, like the young, the elderly, or the infirm. We also expect that the emerging target paradigm could be extended into animal studies in non-human primates and combined with neurophysiological techniques. Together with future work in humans that can rapidly explore the numerous sensory, cognitive, and motor dimensions of the task, the emerging target paradigm should potentiate hypothesis-driven explorations of the fast visuomotor system.

**FIGURE 4.**
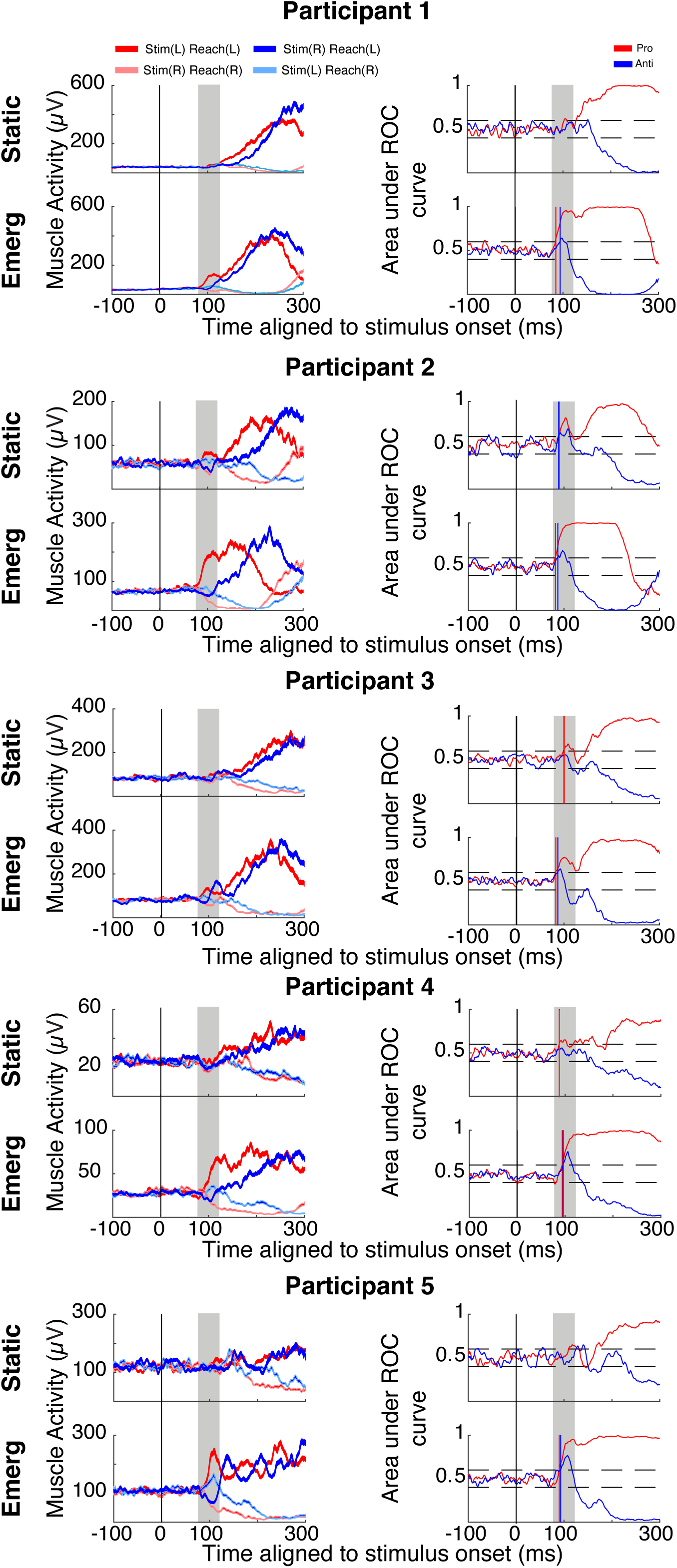
Summary of participant data. Same format as Fig. 2b and 2c, showing data across all five participants.

## ACKNOWLEDGMENTS

This work is supported by a Discovery Grant to BDC from the Natural Sciences and Engineering Research Council of Canada (NSERC; RGPIN 311680) and an Operating Grant to BDC from the Canadian Institutes of Health Research (CIHR; MOP-93796). RAK was supported by an Ontario Graduate Scholarship, and ALC was supported by an NSERC CREATE grant. The experimental apparatus described in this manuscript was supported by the Canada Foundation for Innovation. Additional support came from the Canada First Research Excellence Fund (BrainsCAN).

